# Peak p-values and false discovery rate inference in neuroimaging

**DOI:** 10.1101/358051

**Authors:** Armin Schwartzman, Fabian Telschow

**Affiliations:** Division of Biostatistics, University of California, San Diego

## Abstract

Peaks are a mainstay of neuroimage analysis for reporting localization results. The current peak detection procedure in SPM12 requires a pre-threshold for approximating p-values and a false discovery rate (FDR) nominal level for inference. However, the pre-threshold is an undesirable feature, while the FDR level is meaningless if the signal is assumed to be nonzero everywhere. This article provides: 1) a peak height distribution for smooth Gaussian error fields that does not require a screening pre-threshold; 2) a signal-plus-noise model where FDR of peaks can be controlled and properly interpreted. Matlab code for calculation of p-values using the exact peak height distribution is available as an SPM extension.

## 1 Introduction

Since the introduction of topological inference in neuroimaging (Poline et al., 1997; Chumbley and Friston, 2009; Chumbley et al., 2010), peaks have become a mainstay of neuroimage analysis for localization. Peaks are critical for communicating and reporting neuroimaging results, particularly in task fMRI; tools like BrainMap and Neurosynth.org aggregate peak coordinates studies, providing the basis for meta-analyses. Peaks have also become essential for practical power analysis in neuroimaging (Durnez et al., 2014, 2016). And despite the appeal of cluster inference, peaks remain the most reliable form of topological inference given the inaccuracy of standard Gaussian RFT cluster null distributions (Eklund et al., 2016).

The paradigm for peak inference, proposed by Chumbley et al. (2010) and currently implemented in SPM12, was formally studied in a general setting in Schwartzman et al. (2011) and Cheng and Schwartzman (2017) as the “smoothing and testing of maxima” (STEM) algorithm, and is described there as consisting of the following steps:

1. *Kernel smoothing*: to increase the signal-to-noise ratio (SNR).
2. *Candidate peaks*: find local maxima of the smoothed field above a pre-threshold.
3. *P-values*: computed at each local maximum under the null hypothesis of no signal in a local neighborhood.
4. *Multiple testing*: apply a multiple testing procedure and declare as detected peaks those local maxima whose p-values are significant.

This general recipe relies on two critical elements for proper inference: 1) calculation of valid p-values in Step 3, and 2) interpretation of the nominal error rate in Step 4. The main goal of this article is to formally address these two issues in the context of neuroimaging.

### 1.1 Peak p-values

To calculate p-values, the formula introduced in Chumbley et al. (2010) and the formula currently implemented in SPM12 rely on a screening pre-threshold, so that only peaks higher than the pre-threshold are included in the analysis. SPM users are by now used to the concept of a pre-threshold, as it is routinely used for cluster forming in cluster analysis (Zhang et al., 2009). A pre-threshold has the advantage of reducing the number of features to be tested. However, having to choose a pre-threshold is not necessarily a desirable feature. Analysis results depend on the pre-threshold and change according to the value chosen. This is unsettling for the user. The guidelines for choosing the pre-threshold are loose, only specifying that it should be high enough to enter the regime of the tails of the distribution, but not too high so that it does not eliminate potential discoveries. The dependence of the results on the choice of pre-threshold also opens the door for manipulation of the pre-threshold in order to obtain desired results, which unfortunately invalidates the inference.

The main reason for requiring a pre-threshold for peaks has been one of necessity, as it has been the only known way to obtain approximate p-values. The concept goes back to Adler (1981), who formally defined the *overshoot distribution* as the conditional distribution of peak height above a pre-threshold. Recently, however, an exact formula for the peak height distribution of isotropic Gaussian fields has been obtained (Cheng and Schwartzman, 2015a,b, 2018). When the field has unit variance, this formula relies on a single parameter called *κ*, which depends only on the shape of the autocorrelation function. For example, for an isotropic field with a Gaussian autocorrelation function, this parameter is equal to 1. Otherwise, it can be derived analytically or numerically if the shape of the autocorrelation is known, or otherwise it can be estimated from the data if the field is sufficiently smooth.

Since the peak height distribution depends only on the shape of the autocorrelation function, not its spatial scaling, we conjecture here that its validity is not restricted to isotropic fields but holds more generally for certain classes of nonstationary Gaussian fields, such as anisotropic fields (affine transformations of isotropic fields) and locally stationary fields (where the stationarity property holds locally). In this paper, we evaluate the peak height distribution via simulations for a variety of such fields and test its validity for both Gaussian and *t*-fields. We also propose a method for estimating the required parameter *κ* from data.

More generally, and for fair comparison, we consider the peak p-value formulas from Chumbley et al. (2010) and currently implemented SPM12, formally called approximate peak overshoot distributions as they measure peak heights above a pre-threshold. We compare them with the overshoot distribution from Adler (1981). This distribution was originally derived for stationary fields but was shown by Cheng and Schwartzman (2015a) to be valid for general nonstationary fields. Thus, the use of a pre-threshold is still valuable if the error field is suspected to be highly nonstationary to the extent that the peak height distribution may be invalid. However, this may not be necessary in practice, as shown here in a data example from Moran et al. (2012). Currently there is no formal proof of validity for the overshoot distribution in Chumbley et al. (2010) or SPM12.

### 1.2 False Discovery Rate (FDR) inference

The second critical issue in peak inference is the interpretation of the error rate when declaring statistical significance. Interpretation heavily relies on the definition of what is understood as an error. Chumbley and Friston (2009) make a convincing argument for dismissing the null hypothesis of no signal and argue instead for signal being present everywhere in the brain. This is, in fact, their basis for advocating topological inference based on peaks in Chumbley et al. (2010). However, admitting that the signal is nowhere zero creates a logical fallacy when it comes to statistical testing.

First, p-values are by definition calculated under the null hypothesis. Thus, assuming that the null hypothesis is nowhere true automatically invalidates the calculation of p-values. Second, if the signal is nonzero everywhere, it is impossible to separate between true and false discoveries; every significant peak is a true discovery and the error rate, by definition, is zero. As a consequence, if a multiple testing algorithm is applied with a nominal false discovery rate (FDR) of say, 0.05, one cannot conclude that the FDR is 0.05. The detection algorithm may separate between high peaks that may be considered interesting and low peaks that may be considering less interesting, perhaps in the philosophical style of Efron (2004). This approach was considered by Schwartzman et al. (2009) in the context of voxelwise inference, but it is more difficult to formalize and implement in the context of peak inference and we do not pursue it here.

The only way to achieve meaningful FDR inference in the traditional statistical sense and commonly understood by most investigators is to allow the signal to be zero in some regions and nonzero in others. Even Chumbley and Friston (2009) themselves admit that, although smoothing with a Gaussian kernel in theory creates signal everywhere, the kernel in practice has finite support and does not produce that effect. Cheng and Schwartzman (2017) solve the problem assuming a model where the signal is nonzero only on a set of unimodal functions with finite support. Being unimodal, each signal component is topologically represented by its own mode. This model allows a precise and straightforward definition of statistical inference for peaks: a significant peak that falls within the support of a signal component is a true discovery, while a significant peak that falls in the zero-signal region is a false discovery. Given these definitions, the FDR can be appropriately defined as the expected proportion of falsely discovered peaks. We define the detection power as the expected fraction of truly discovered peaks.

In Cheng and Schwartzman (2017) it is shown that, if the noise field is stationary and ergodic, then the FDR of peaks is asymptotically controlled as both the sample size and the search domain increase. It is also shown that if the exact height distribution is used, then pre-thresholding reduces detection power and it is best not to apply pre-thresholding at all. This is contingent on the height distribution being correctly specified. If the approximate overshoot distribution is used instead, then pre-threshold can be optimized to balance accuracy against detection power. The simulation examples in Cheng and Schwartzman (2017) were limited to 2D fields. In this paper, we perform simulations to evaluate FDR control and detection power in 3D fields, more representative of neuroimaging, and under a variety of noise field conditions.

We emphasize that the purpose of the above argument is not to invalidate the scientific premise of Chumbley and Friston (2009) that there may be signal everywhere in the brain, and surely not the neurological arguments supporting it. Our goal is only to provide a framework where statistical inference for peaks may be taken to be valid and properly interpreted.

The rest of the paper addresses the calculation of p-values for peaks in Section 2 and the FDR inference in Section 3. The specific implementation for fMRI is discussed in Section 4 and some concluding remarks are provided in Section 5. Matlab code for calculation of p-values using the exact peak height distribution is available in GitHub (https://github.com/fjete88/STEM) and as an SPM extension in the SPM12 Extensions website (http://www.fil.ion.ucl.ac.uk/spm/ext//#STEM).

## 2 Peak p-values

### 2.1 Peak height distribution

Suppose that *z*(*s*) is a smooth Gaussian random field, with mean 0 and variance 1, indexed by a continuous space location index *s* in a Euclidean space of dimension *D*. Smoothness here refers to the field being 3-times differentiable^1^. This holds if the smoothing kernel is itself 3-times differentiable, such as a Gaussian or a quartic kernel.

The peak height distribution of a local maximum of *z*(*s*), as a function of a height threshold *u*, is the probability that *z*(*s*) is greater than *u* given that a local maximum has occurred at *s*. This is *not* the same as the marginal probability

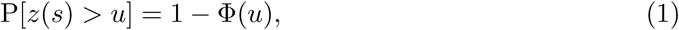

where Φ(*u*) is the cumulative distribution function (cdf) of a standard normal random variable. Marginal peak p-values using (1) are too liberal because they do not take into account the fact that peaks have been selected for being local maxima. Peaks are, by definition, higher than their neighboring voxels, and therefore are more likely to be high than any voxel chosen at random. This distinction is important particularly when reading the output of SPM12, which still reports marginal p-values for peaks; those p-values are misleadingly too low.

The peak height distribution is properly defined as the *conditional* probability

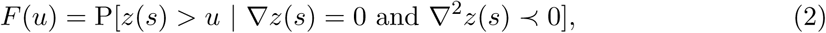

where *∇ z*(*s*) is the gradient of *z* at *s*, *∇*^2^*z*(*s*) is the Hessian of *z* at *s*, and the symbol *≺* indicates negative definiteness, specifying *s* as a local maximum. The conditioning terms in (2) specify the selection criteria and make this distribution stochastically greater than the standard Gaussian distribution (1). Figure 1 illustrates the difference between the two distributions in the case of an isotropic field. Note that the peak distribution is always shifted toward higher values with respect to the standard normal.

**Figure 1.**
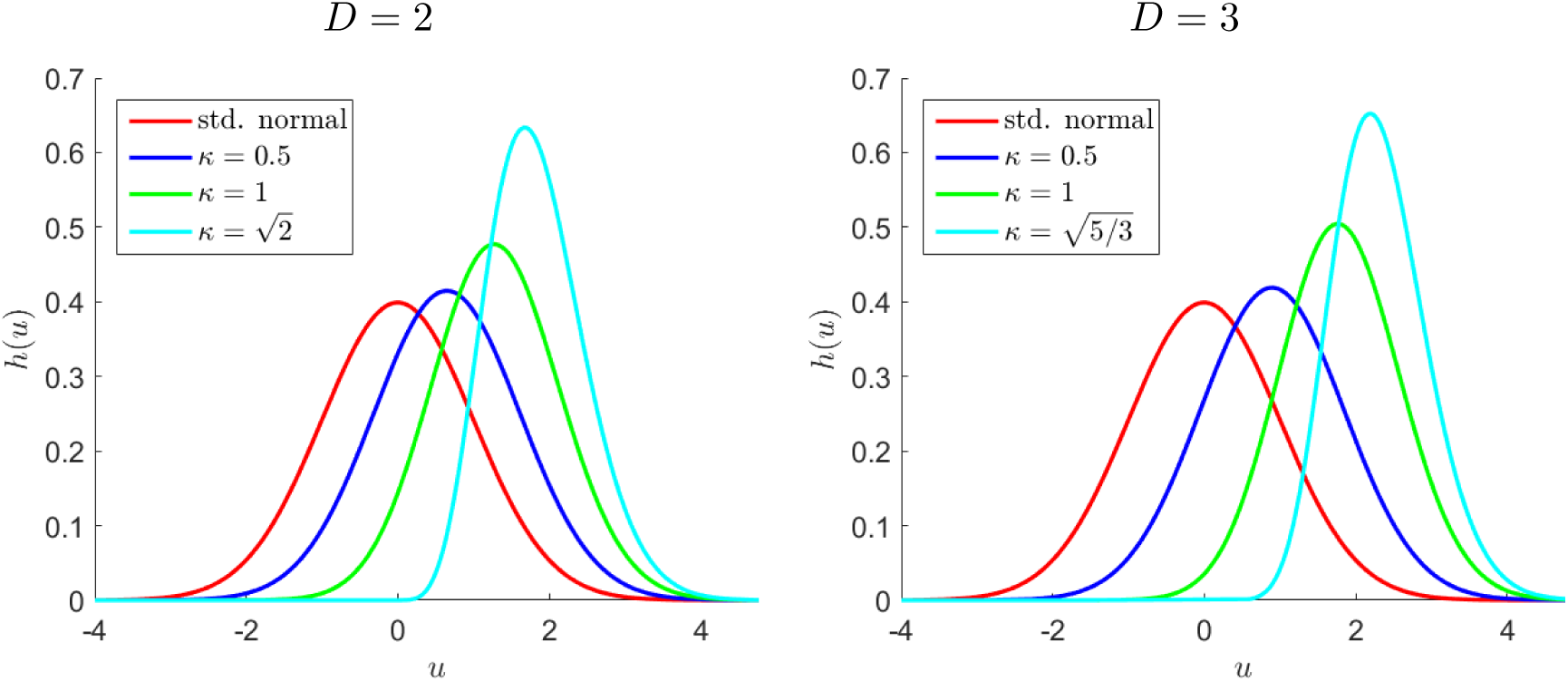
Density functions *h*(*x*; *κ*) of the height distribution of local maxima for dimensions *D* = 2 (left) and *D* = 3 (right). The largest value of *κ* allowed is 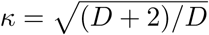

### 2.2 Peak height distribution for isotropic Gaussian fields

When the field *z*(*s*) is isotropic, i.e. its covariance function is invariant under translations and rotations, then the height distribution of local maxima has a closed-form expression for dimension up to *D* = 3 (Cheng and Schwartzman, 2018, 2015b). Specifically, the tail probability (2) can be written as

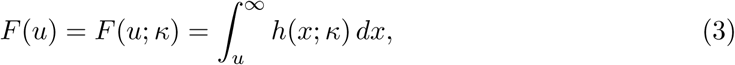

where *h*(*x*; *κ*) is the probability density function (pdf) of the height of local maxima, depending on a single parameter *κ*. When *D* = 2, the pdf is given by

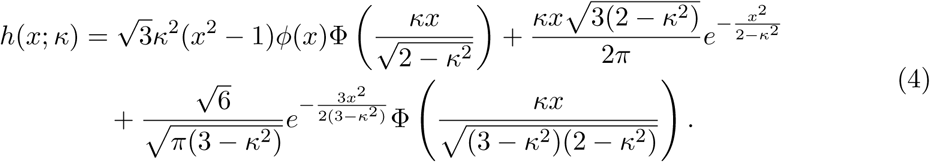

When *D* = 3, the pdf is given by

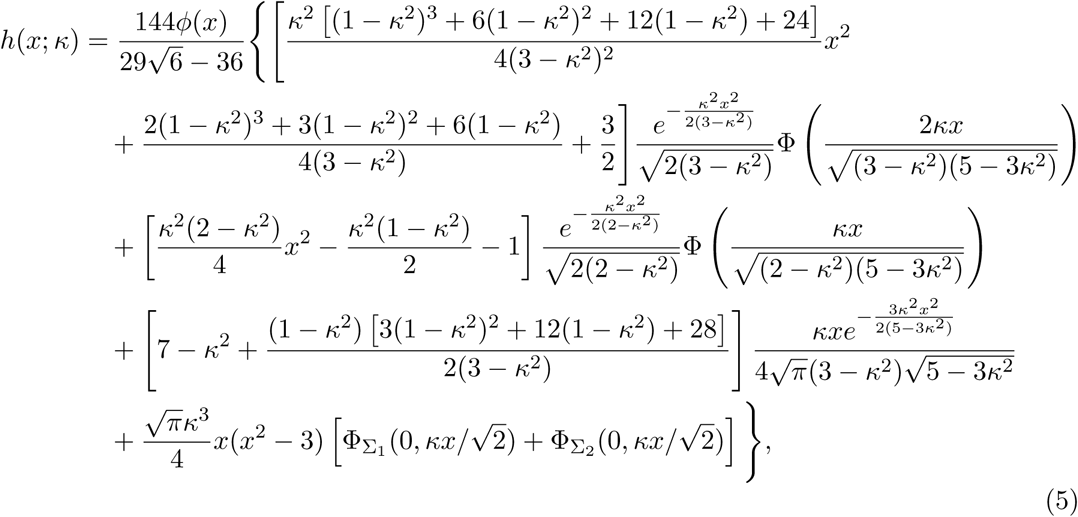

where Φ_Σ1_ and Φ_Σ2_ denote bivariate normal cdfs corresponding to the covariance matrices

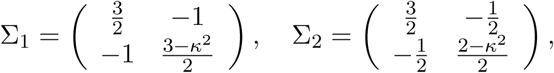

respectively. The densities (4) and (5) for various values of *κ* are plotted in Figure 1. Note that increasing *κ* shifts the distribution toward higher values.

In the densities (4) and (5), the single parameter *κ* contains all the necessary information about the covariance function of the field and is defined as follows. Recall that *z*(*s*) is assumed to have mean 0 and variance 1. If we write the autocorrelation function (acf) of the field as 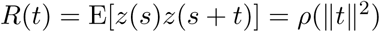 for an appropriate function *ρ*, then

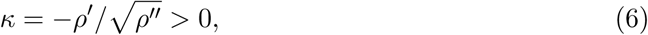

where 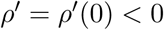 and 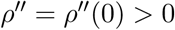 are the first and second derivatives of the function *ρ* evaluated at zero (Cheng and Schwartzman, 2015a). Common examples are the following:

- *Squared exponential or Gaussian acf*: Here 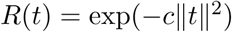 with 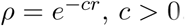. Thus 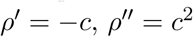 and *κ* = 1.
- *Inverse squa redpolynomial or Cauchy acf*: Here 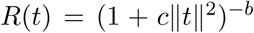 with *ρ(r)* = (1 + *cr*)^−*b*^, *c* > 0, *b* > 0. Thus 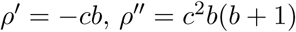, and 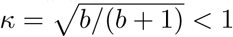

In general, the largest value that *κ* may take is 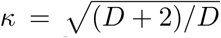, obtained in the special case of a field satisfying Helmholtz’s equation 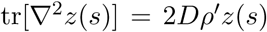 (Cheng and Schwartzman, 2018).

An interesting property of the parameter *κ*, following directly from definition (6), is that it is invariant under scalings of the space. That is, *κ* is the same for the field *z*(*s*) as for the field *z*(*as*), for any fixed real number *a*. Because of this property, the values of *κ* in the examples above do not depend on the constant *c*. More importantly, it implies that the peak height distribution is scale invariant as well and depends only on the shape of the acf near the origin. This is not surprising, as scaling of the space does not affect the values of the field themselves. We exploit this scale-invariant property in the simulations below.

### 2.3 Simulations of isotropic Gaussian fields

To confirm formula (5) in a finite resolution setting, we simulated 1000 random fields of size 50 × 50 × 30 voxels, each field obtained as the convolution of white Gaussian noise with an isotropic kernel *w*(*s*), properly accounting for the margins to ensure validity of the convolution. The generated fields were normalized by the factor 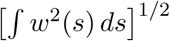 to ensure unit variance.

Figure 2 shows the results using a Gaussian kernel with standard deviation *ς*, which we shall call bandwidth (it relates to the full-width half max (FWHM) via FWHM *≈* 2.355 *ς*). The figure shows an example slice of the simulated field, the histogram of the measured peak heights compared to the theoretical density (5) with *κ* = 1, and the empirical distribution of the corresponding p-values. As shown in the figure, the accuracy of the formula improves with increasing smoothness. As the bandwidth increases, the empirical peak height distribution becomes closer to the theory and the empirical p-value distribution becomes closer to the uniform distribution. On the other end, when the smoothing kernel is not large enough (e.g. *ς* = 3), the height distribution is stochastically smaller than the theory. This is because the true peaks fall between the grid points and thus the measured peak heights are slightly smaller than they should be. Fortunately, this implies that the p-value distribution is stochastically larger than the uniform. Larger p-values are still valid for inference, albeit conservative.

**Figure 2.**
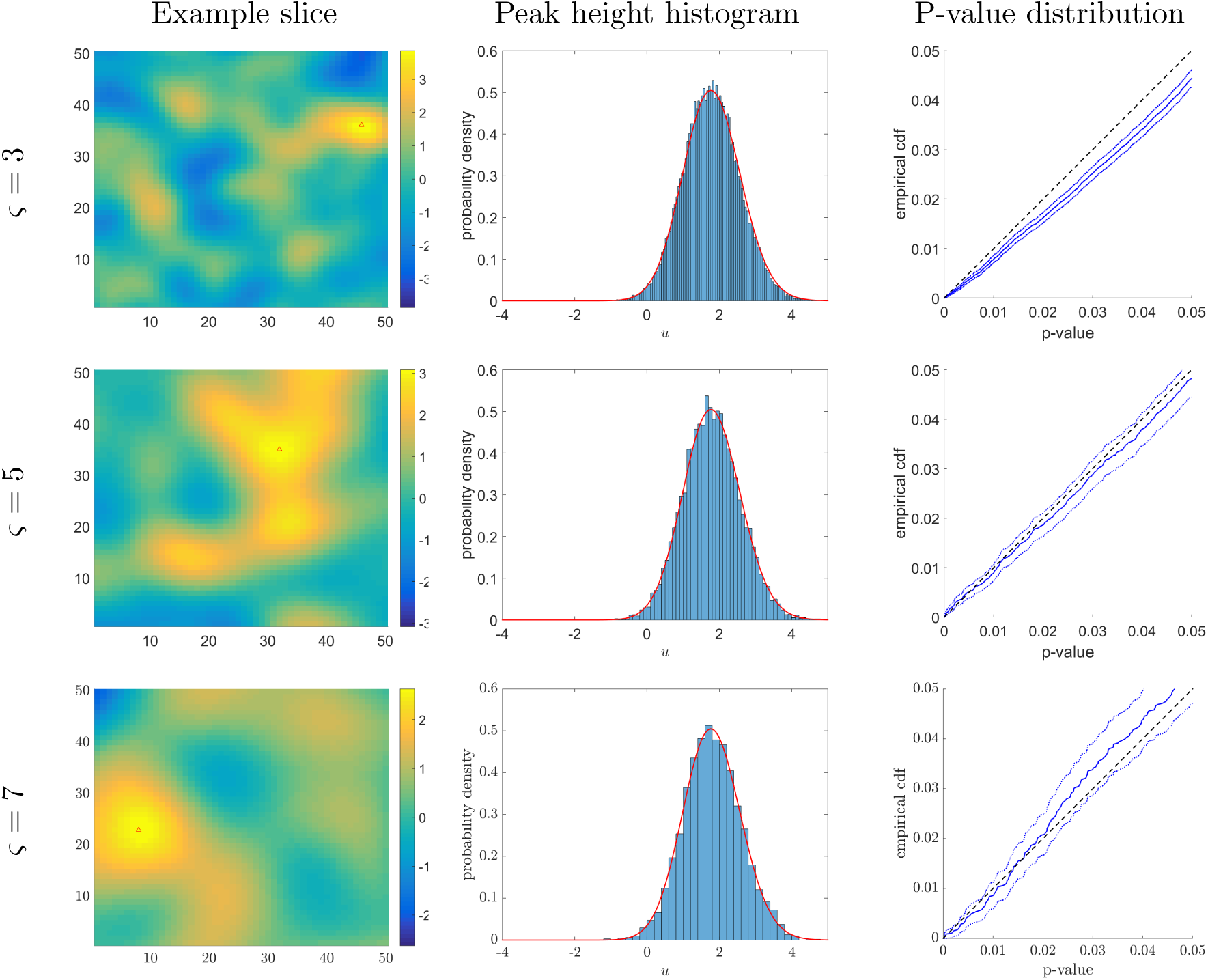
Peak height simulations for an isotropic Gaussian field generated with a Gaussian kernel. *Left column:* The slice containing the global maximum (marked with a triangle). *Middle column:* Histogram of peak heights over 10,000 simulated fields; superimposed is the theoretical peak height density with *κ* = 1. *Right column:* Empirical p-value distribution with confidence envelope.

To confirm formula (5) in the case of a non-Gaussian autocorrelation function, we repeated the above simulations (1000 random fields of size 50 × 50 × 30 voxels), using instead an isotropic 3D quartic (biweight) kernel

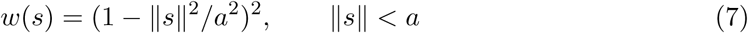

with *a* = 18 voxels (corresponding to about 2.5 bandwidths of a Gaussian with *ς* = 7, to approximately match the smoothness in the last row of Figure 2). The value of *κ* for the isotropic quartic kernel was computed numerically as *κ* = 0.92 (see Appendix A.1). Figure 3 shows the same panels as Figure 2, confirming the validity of the distribution for this kernel.

**Figure 3.**
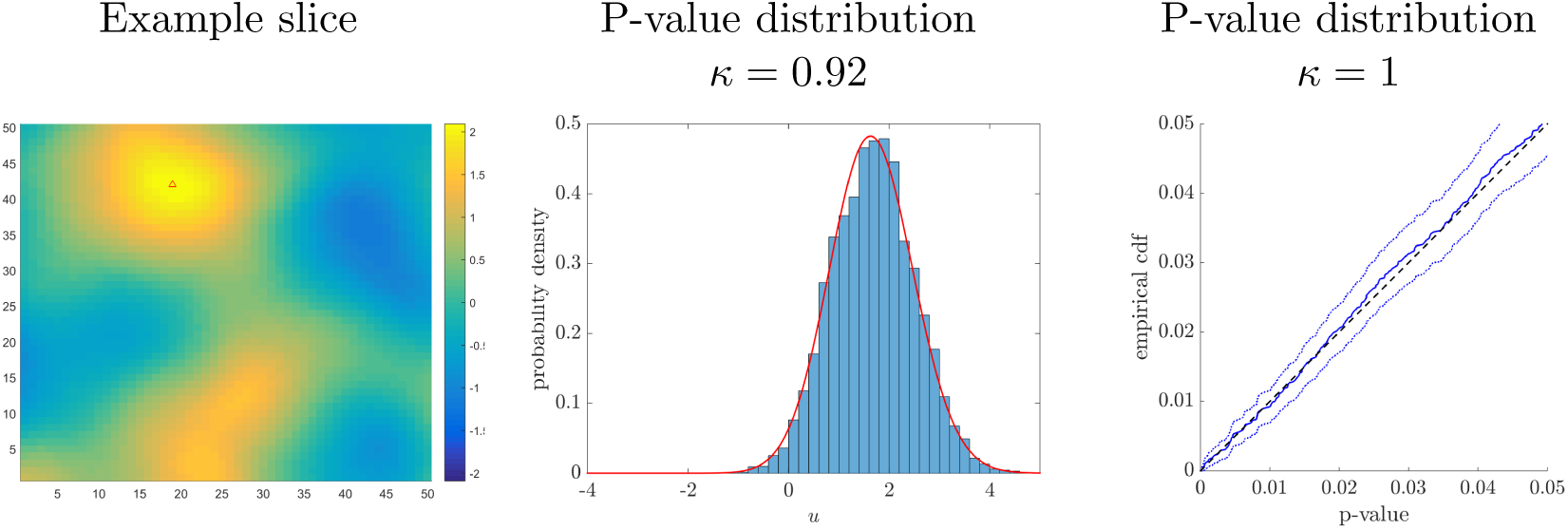
Peak height simulations for an isotropic Gaussian field generated with a quartic kernel. *Left column:* The slice containing the global maximum (marked with a triangle). Middle: Histogram of peak heights over 10,000 simulated fields; superimposed is the theoretical peak height density with *κ* = 0.92. *Right column:* Empirical p-value distribution with confidence envelope.

### 2.4 Unknown *κ*

In the simulations above, the value of *κ* was assumed to be known. This is generally not the case in practice. The parameter *κ* can be estimated from the data as follows. Suppose that the observed field is 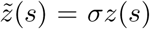 with mean 0 and variance *σ*^2^. Following definition (6), it is shown by Cheng and Schwartzman (2017) that

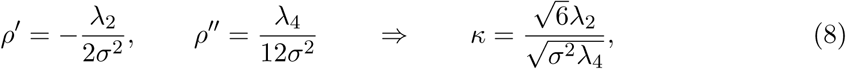

where *λ*_2_ and *λ*_4_ are the spectral moments of the field. These are defined as the variance of the first and second spatial derivatives of the field 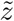, respectively, at any location and in any direction in space (assuming that the field is isotropic). The variances *σ*^2^, *λ*_2_ and *λ*_4_ can be estimated empirically from the numerical derivatives in all three spatial directions and at all spatial locations. If multiple instances of the field are available, then they are all included in the calculation of the variances. Plugging in (8) yields an estimate of *κ*.

Table 1 shows the estimation accuracy of the above method when the field is produced as a convolution of white Gaussian noise with a Gaussian or a quartic kernel. As the size of the Gaussian smoothing kernel increases, the estimated value gets closer to the theoretical ideal value of *κ* = 1. For the quartic kernel, the estimated value is slightly higher than the theoretical value *κ* = 0.92 because of the limited spatial resolution. For both kernels, the discreteness of the grid biases the numerical derivatives and results in a small bias in the estimation of *κ*. In all cases, the standard error decreases with the sample size, as expected.

**Table 1.**
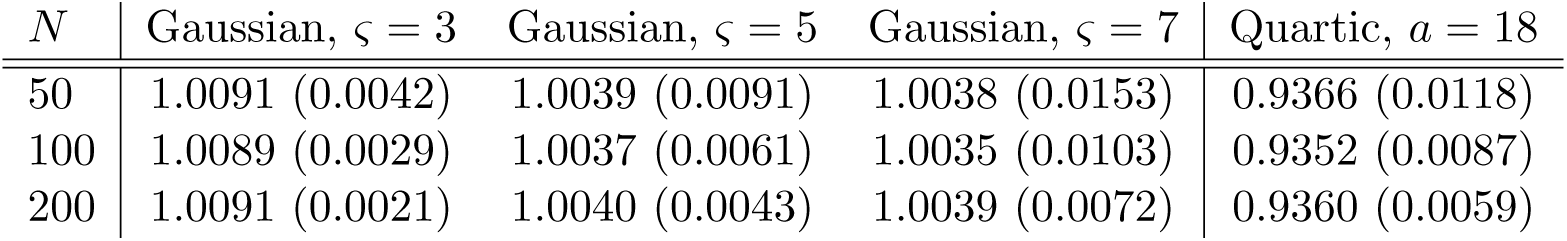
Estimation of the parameter *κ* as a function of sample size *N* for various smoothing kernels. Simulation standard errors are in parentheses.

### 2.5 Simulations of non-isotropic fields

The scale-invariant property indicates that the peak height distribution only depends on the shape of the covariance function. This suggests that the height distribution may still be valid if the scaling factor is different in every direction in space, i.e. if the field is anisotropic, or if it changes slowly over space, i.e. if the field is mildly nonstationary.

To evaluate these claims, we performed simulations as above (1000 random fields of size 50 × 50 × 30 voxels) under an anisotropic scenario and a nonstationary scenario. In the anisotropic scenario, the kernel was Gaussian anisotropic with bandwidths *ς* = 5, 7, and 9 in the three spatial directions. In the nonstationary scenario, half of the white noise field (before smoothing) was binned into blocks of 2 × 2 voxels. Then the entire noise field was convolved with a Gaussian kernel with bandwidth *ς* = 7, producing a smooth field with effectively a different acf in the two halves. The fields were normalized to have unit variance at every voxel.

Figure 4 shows the peak height and p-value distributions. The peak height distribution appears to be valid in both these scenarios.

**Figure 4.**
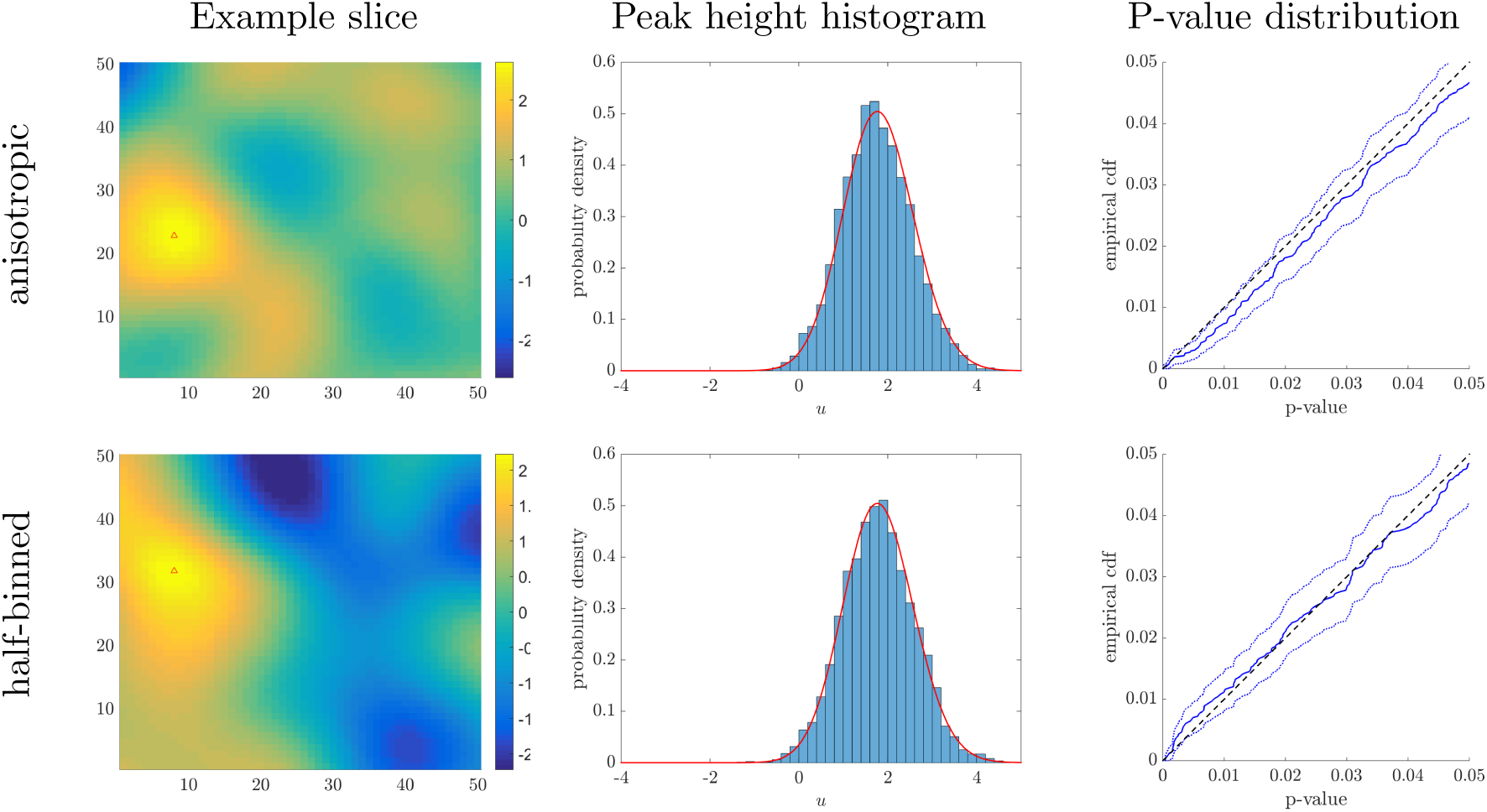
Peak height simulations for an anisotropic Gaussian field (top row) and half-binned nonstationary field (bottom row), generated with a Gaussian kernel. *Left column:* The slice containing the global maximum (marked with a triangle). *Middle column:* Histogram of peak heights over 10,000 simulated fields; superimposed is the theoretical peak height density with *κ* = 1. *Right column:* Empirical p-value distribution with confidence envelope.

### 2.6 Application to *t*-fields

In practice, the null distribution of the test statistic map is typically *t* rather than Gaussian. In Figure 5, *t*-fields where created by voxelwise calculation of a *t*-statistic using

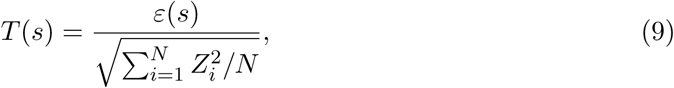

where *Z*_1_,*…, Z _N_* and *ε*(*s*) are i.i.d. isotropic Gaussian fields such as those simulated in Figure 2. Because the exact height distribution is designed for Gaussian fields, not *t*-fields, the p-value distribution in Figure 5(left) is too liberal, although it improves as the number of d.f. increases.

**Figure 5.**
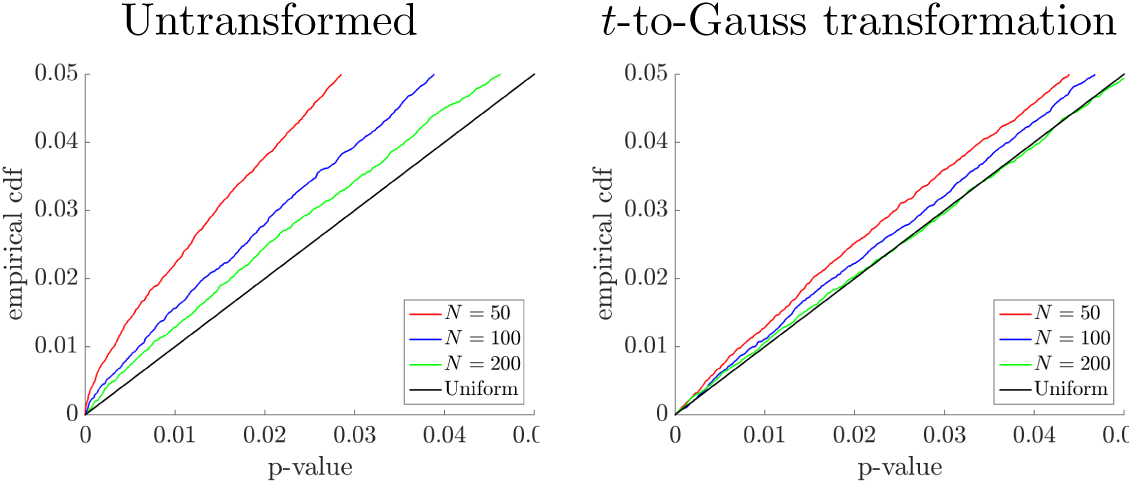
Peak height simulations for a *t* field with *N* d.f. (10,000 simulations). *Left:* Direct application of the exact Gaussian height distribution. *Right:* Exact Gaussian height distribution after a *t*-to-Gaussian quantile transformation.

To improve the accuracy for *t*-fields, we transform the *t*-fields to be marginally Gaussian by means of the quantile transformation

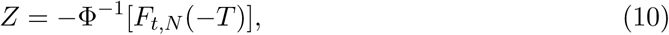

where *F _t,N_* is the cdf of the *t* distribution with *N* d.f. The minus signs ensure the accuracy of the transformation at the upper tail of the distribution, where it matters most. We then apply the exact Gaussian height distribution to the transformed *t*-fields. Figure 5(right) shows that the p-value distribution is now very accurate, especially if the number of d.f. is higher than 100.

### 2.7 Overshoot distribution

Rather than the height distribution of local maxima, the peak inference method in Chumbley et al. (2010) and the method currently implemented in SPM use the overshoot distribution. The overshoot distribution (Adler, 1981) is defined as the probability that a peak is higher than *u* given that it is already higher than a pre-threshold *v < u*:

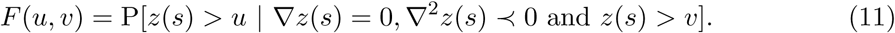

Using conditional probability rules, the overshoot distribution (11) can be readily obtained as

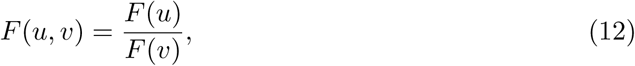

where *F*(*u*) is the peak height distribution (2). For isotropic Gaussian fields, the overshoot distribution can be calculated exactly replacing *F*(*u*) = *F*(*u*; *κ*) as given by (3).

For smooth non-isotropic Gaussian fields, various approximations to the overshoot distribution have been offered in the past. All the approximations become more accurate as the pre-threshold *v* increases. The earliest one is Adler’s approximation (Adler, 1981; Adler et al., 2010)

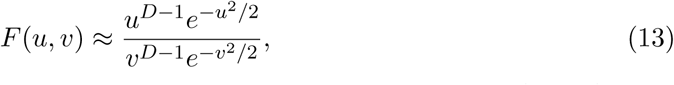

originally derived for stationary fields but shown in Cheng and Schwartzman (2015a) to be asymptotically valid also for nonstationary fields.

For Gaussian fields, Chumbley et al. (2010) proposed the approximation

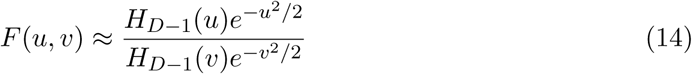

in the Gaussian case, where *H_D−_*_1_(*x*) is the Hermite polynomial of order *D* − 1. This approximation was also derived independently by Cheng and Schwartzman (2015a) using the Kac-Rice formula and was shown rigorously there to be accurate for stationary smooth Gaussian fields. Note that (14) is the same as (13) when *D* = 2. For a *t*-field with *ν* degrees of freedom, Chumbley et al. (2010) proposed the analogous approximation

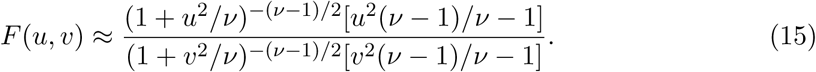

In SPM12, peak p-values are calculated according to the approximation

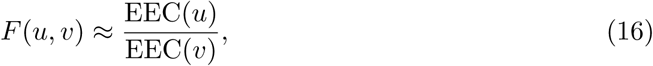

where EEC(*u*) is the expected Euler characteristic at level *u*, given in its usual form by the Gaussian kinematic formula using random field theory for Gaussian or *t*-fields (Worsley et al., 1996, 2004). Note that this approximation reduces to (14) in the Gaussian case or (15) in the *t* case if only the highest order term is taken in the expansion of the EEC.

Figure 6 compares the various approximations above in the Gaussian isotropic case for a fixed pre-threshold of *v* = 2.5. For *D* = 2, the Chumbley et al. (2010) and Adler (1981) approximations are the same. For *D* = 3, the Adler (1981) approximation (13) is almost exact. In general, the Chumbley et al. (2010) approximation (14) is conservative and the SPM12 approximation (16) is liberal, although the effect is most noticeable for *D* = 3.

Note that both the exact overshoot approximation (12) and the SPM12 approximation (16) require estimation of parameters, *κ* in the former and the Lipschitz-Killing curvatures (LKCs), or resel counts, in the latter. To separate the estimation problem from the form of the overshoot approximation, we use the exact theoretical values of these parameters in the simulations to follow. In addition, estimation of the LKCs is slow in SPM12; avoiding it results in valuable savings in computation time over multiple repeated simulations.

### 2.8 Overshoot distribution simulations

To compare the various options for the overshoot distribution, we performed simulations as above (random fields of size 50 × 50 × 30 voxels) under an isotropic scenario.

The first set of simulations evaluates the effect of the pre-threshold. Here, white Gaussian noise was convolved with an isotropic Gaussian kernel with bandwidth *ς* = 7, as in Figure 2 (bottom row). Figure 7 shows that the exact distribution (12) is precise for all values of the pre-threshold *v*, while the other approximations get better as *v* increases. Consistent with Figure 6, the Chumbley approximation (14) in Figure 7 gives conservative p-values (stochastically larger than uniform) while the SPM12 approximation (16) gives liberal p-values. In other words, the Chumbley p-values are valid, albeit conservative, but the SPM12 p-values are not. We will see in Section 3.3 below that SPM12 yields a higher rate of false positives in peak detection.

**Figure 6.**
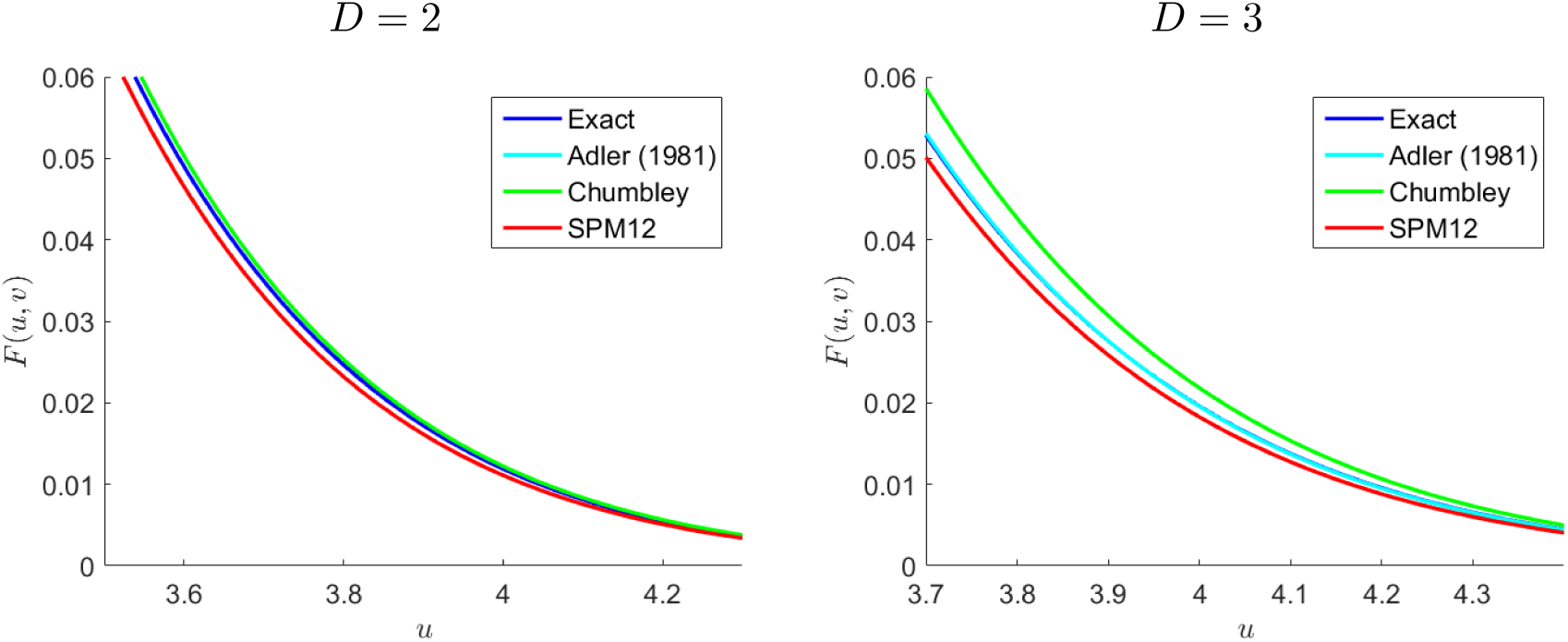
Overshoot probabilities *F*(*u, v*) for dimensions *D* = 2 (left) and *D* = 3 (right), pre-threshold *v* = 2.5.

**Figure 7.**
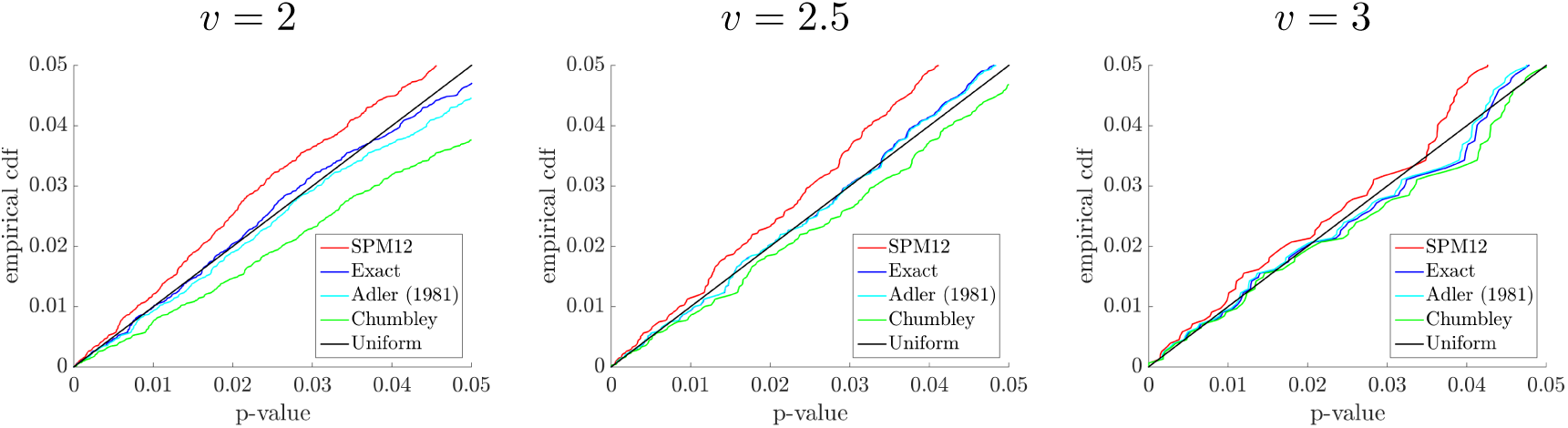
Overshoot p-value distributions for an isotropic Gaussian field obtained with a Gaussian kernel with bandwidth *ς* = 7 (10,000 simulations), as a function of the pre-threshold *v*.

Figure 8 (top row) compares the overshoot distribution approximations for *t*-fields with *N* d.f. for a fixed pre-threshold *v* = 2.5. Here, *t*-fields where created by voxelwise calculation of a *t*-statistic from isotropic Gaussian fields as those simulated in Figure 7. The exact distribution (12) is inaccurate because it is designed for Gaussian fields, but improves as the number of d.f. increases. The Chumbley et al. (2010) approximation for *t*-fields is accurate for any number of d.f. Surprisingly, the SPM12 approximation for *t*-fields is biased and does not improve with increasing d.f.

**Figure 8.**
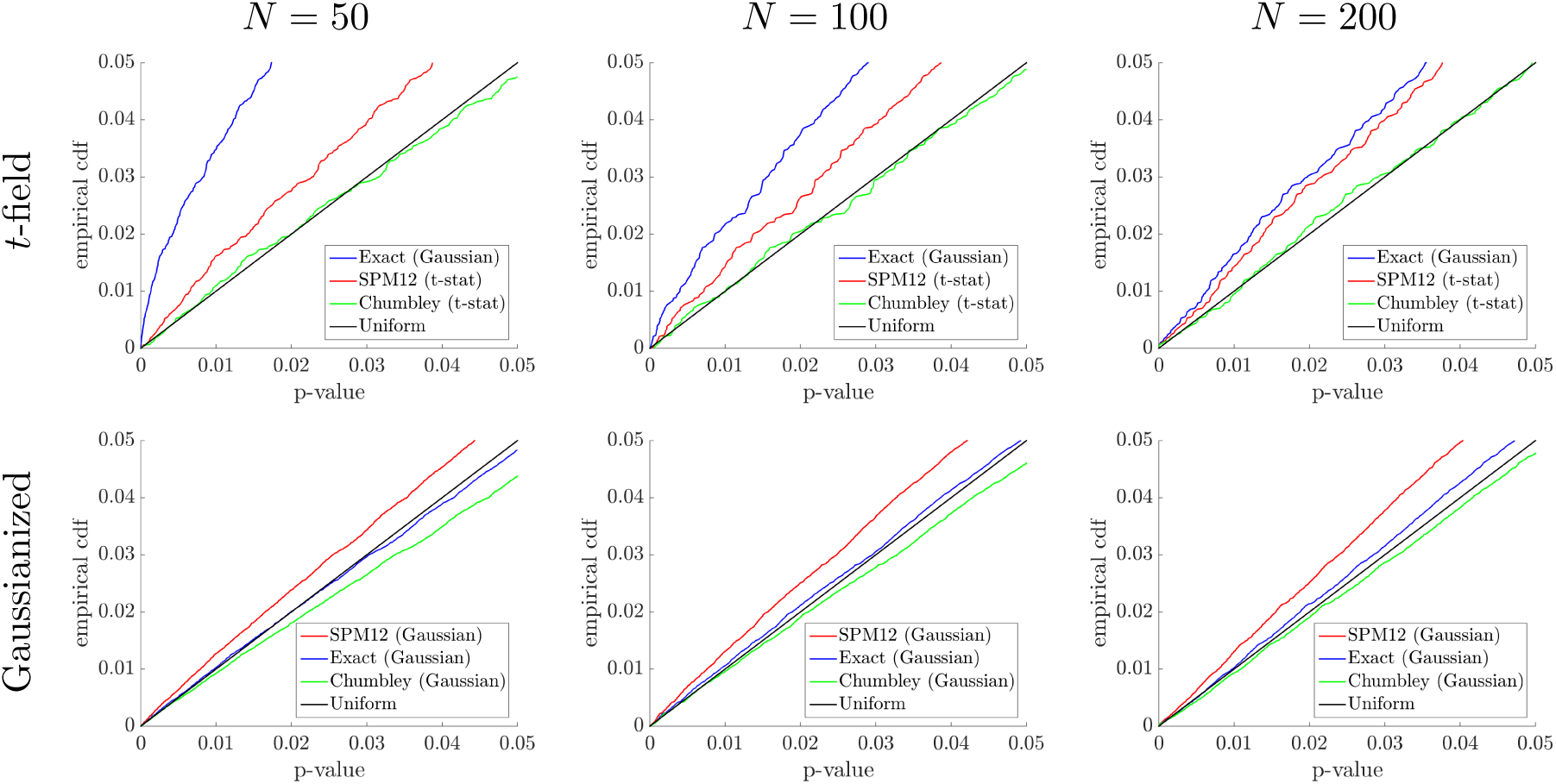
*Top row:* Overshoot p-value distributions for an isotropic *t*-field with *N*d.f. (10,000 simulations), generated from isotropic Gaussian fields (Gaussian kernel with bandwidth *ς* = 7). The pre-threshold is fixed at *v* = 2.5. *Bottom row:* Same as top row, after *t*-field is transformed by a *t*-to-Gaussian quantile transformation (100,000 simulations).

In contrast, Figure 8 (bottom row) shows the results when the *t*-fields are transformed to marginally Gaussian by the quantile transformation (10). Here we used 100,000 simulations to better discriminate between the methods. Applied to the Gaussianized fields, the exact overshoot distribution is very accurate. On the other hand, the Gaussian version of Chumbley’s approximation is slightly conservative, while the Gaussian version of the SPM12 approximation remains biased in the liberal direction.

## 3 FDR inference

In proposing FDR inference for peaks, Chumbley et al. (2010) not only argued for peaks being topological quantities of interest, but also argued against the null hypothesis of no activation. In their view, activity is pervasive in the brain; the null hypothesis of no activation is never true and can always be rejected with a large enough sample size. We do not necessarily argue against the scientific validity of this claim, but against its statistical interpretation. The problem with this view is that without a null hypothesis to reject, there is no meaning to the concepts of statistical testing and error rates. Inference as desired by Chumbley et al. (2010) is not well defined and nominal error rates cannot be interpreted as such.

To make peak inference meaningful, Schwartzman et al. (2011) in the one-dimensional case, and later Cheng and Schwartzman (2017) in the *D*-dimensional case, provided a setting under which peak detection could be made rigorous and FDR could be properly interpreted. The detection algorithm is essentially the same as that of Chumbley et al. (2010), but the interpretation is different.

### 3.1 Modeling assumptions

To control FDR in a rigorous manner, Cheng and Schwartzman (2017) show that this is possible under certain modeling assumptions. Cheng and Schwartzman (2017) work under a signal+noise model

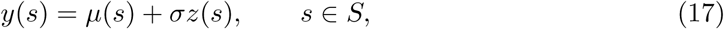

where *µ*(*s*) represents the signal and *σz*(*s*) the noise with *σ >* 0, both defined on a search region *S ⊂ ℝ^D^*. As shown in Section 4 below, coefficient estimates from the general linear model in fMRI can be written in this form.

The signal *µ*(*s*) itself is modeled as a linear composition of *J* bump functions

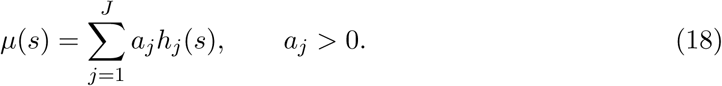

Each bump function *h_j_*(*s*) is positive, unimodal and has compact connected support, representing a peak to be detected. For identifiability of the coefficient *a _j_*, each bump function *h_j_*(*s*) is required to have unit action, i.e. integral equal to 1. In addition, bump functions are required to be twice differentiable within their support and have no critical points other than their mode. These functions need not be concave within their support nor their level sets need to be convex, allowing for a large variety of possibilities for the signal *µ*(*s*).

For the noise term *z*(*s*), in addition to it being a smooth Gaussian field as in Section 2, here it is required that the noise field be stationary ergodic. Ergodicity refers to the property that statistics of the field over many realizations (e.g. moments or excursion probabilities) can be estimated from a single realization of the field by empirical calculations over space if the domain is large enough. Ergodicity usually holds if the autocorrelation function of the field decays fast enough, that is, if the values of the field at different locations tend to be less correlated as the points are taken farther apart from each other. It does not hold if correlation reaches far in space, for example, if the noise is a periodic function over space.

### 3.2 FDR control

Because the location of truly detected peaks may shift as a result of noise, a significant local maximum is called a true positive in Cheng and Schwartzman (2017) if it falls anywhere inside the support of a true peak; otherwise, it is called a false positive. This allows for a formal definition of FDR as the proportion of falsely detected peaks among significant peaks. It also allows for a formal definition of detection power as the expected fraction of truly detected peaks out of all the *J* existing peaks. This detection power turns out to be the same as the average probability of detection for each of the existing peaks.

Proceeding as in Chumbley et al. (2010), after kernel smoothing of the data, local maxima are found, a p-value is attached to each one of them, and an FDR-controlling procedure is applied to the list of peak p-values to determine a significance threshold.

When applying the BH procedure to the list of peak p-values, Cheng and Schwartzman (2017) formally show that:

1. The FDR tends to *αM/*(*M*+ *J*), where *J* is the number of true peaks and *M* is the expected number of local maxima over the null region.
2. The detection power tends to 1.

Both these results hold asymptotically as the search space and the signal-to-noise ratio (SNR) increase, with the SNR being required to increase faster than the logarithm of the size of the search space. In other words, the size of the search space may be almost exponentially larger than the SNR. This type of scaling relationship is typical of high dimensional data problems (e.g. Candes and Tao (2007); Zhang (2010)).

The reasons for requiring both the search space and the SNR to increase are twofold. First, increasing search space is required to apply ergodicity. As the search space increases, more peaks appear, both true and false. By ergodicity, the empirical distributions of both true and false peak heights involved in the BH procedure converge to the theoretical distributions, allowing nominal FDR control.

Second, increasing SNR is necessary to avoid error inflation from smoothing and to guarantee power consistency. As rightly pointed out by Chumbley and Friston (2009) and Chumbley et al. (2010), convolution with a kernel artificially extends the signal beyond its original domain. This additional extent is called *transition region* in Cheng and Schwartzman (2017). Different solutions to this problem have been offered in the literature. For example, Pacifico et al. (2007) suggested “shaving” the extra signal, although they found their procedure difficult to calibrate so that it would not shave too much. In Cheng and Schwartzman (2017), the assumptions of unimodality and lack of other critical points for the peak functions in (18) guarantee that, as the SNR increases, the probability of observing any peaks in the transition region goes to 0. In other words, asymptotically, peaks occur either in the null region where the signal is zero, or very close to true peaks, eliminating the error inflation. As an additional benefit, the detection power goes to 1.

The expected number of local maxima over the null region required in the limit of the FDR above can be calculated as (Cheng and Schwartzman, 2015b)

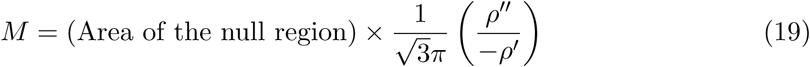

for *D* = 2 and

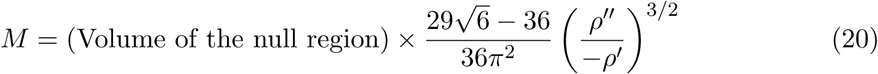

for *D* = 3, where *ρ*′ and *ρ*″ are the same quantities used in the calculation of *κ* in (6). Formulas (19) and (20) do not depend on the variance of the field and remain the same if the noise field does not have variance 1.

### 3.3 Simulations of FDR control and detection power

The simulations in Cheng and Schwartzman (2017) were limited to two dimensions. To evaluate the methods in a situation closer to neuroimaging, we present here simulation results in a 3D setting. As signal, three non-overlapping signal bump functions were placed in a volume of size 50 × 50 × 30 voxels. The signal bumps were generated as 3D outer products of 1D quartic kernels with varying support sizes, heights within 25% of each other, and scaled by a global constant *A* to control the SNR (Figure 9).

**Figure 9.**
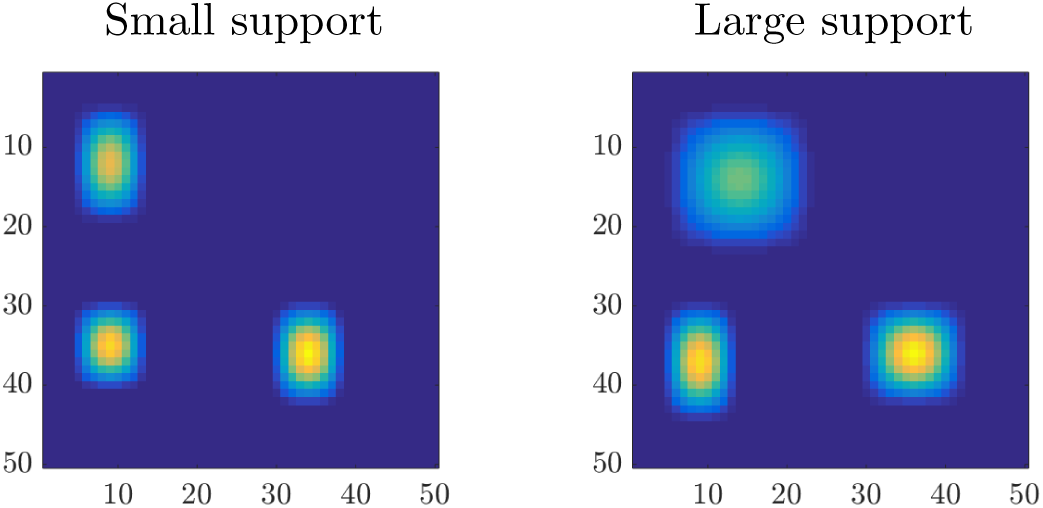
Projection of the signal along the axis perpendicular to the page; in reality the three bumps belong to different slices. Two simulation scenarios are shown with small and large support.

The first simulation setup considers Gaussian noise. Here the noise field was produced by convolution of white noise with a Gaussian kernel with bandwidth *ς* = 5 or *ς* = 7 using the SPM12 function `spm_smooth` and renormalizing to unit variance. The SNR is controlled by scaling the bumps so that the ratio (height of second highest peak) / (standard deviation of noise field) ranges from 3 to 7.

Figure 10 (left column) shows the realized FDR and detection power as a function of the SNR for a nominal FDR level *α* = 0.05 when using the exact height distribution. As predicted by the theory, as the SNR increases, the FDR converges to *αM/*(*M*+ *J*) and the detection power increases to 1. The difference between the two asymptotic FDR values is due to the noise acf. A larger noise bandwidth implies a smaller number of noise peaks *M*, and thus a smaller fraction *αM/*(*M*+ *J*). Here *J* = 3 and *M* was calculated using (20).

**Figure 10.**
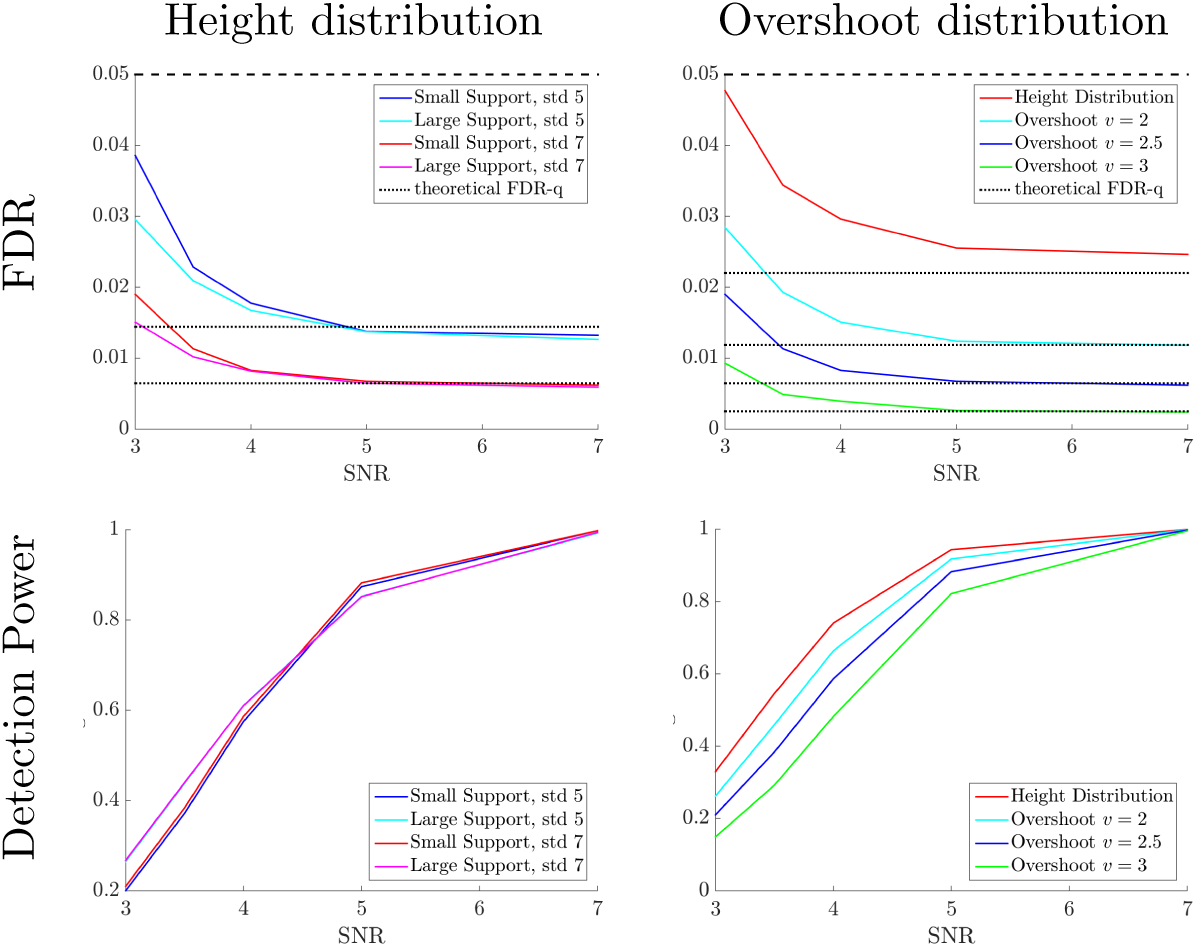
Realized FDR and average detection power (10,000 simulations). Simulated fields are isotropic Gaussian (Gaussian kernel with bandwidth *ς* = 7).

Figure 10 (right column) compares the realized FDR and detection power when using the exact height distribution versus the exact overshoot distribution in the calculation of peak p-values. Increasing the screening pre-threshold results in a smaller number of noise peaks and thus a smaller limiting FDR value, but also a smaller detection power. Using the exact height distribution with no pre-threshold results in the highest detection power while still keeping the FDR below the nominal level.

Figure 11 compares the various approximations to the overshoot p-value distribution in terms of the realized FDR and detection power. As the pre-threshold increases, the theoretical FDR limit decreases because the expected number of local maxima in the null region decreases. Fortunately, there is enough margin so that, even if the p-value distributions are only approximate, the FDR is still below the nominal level 0.05.

**Figure 11.**
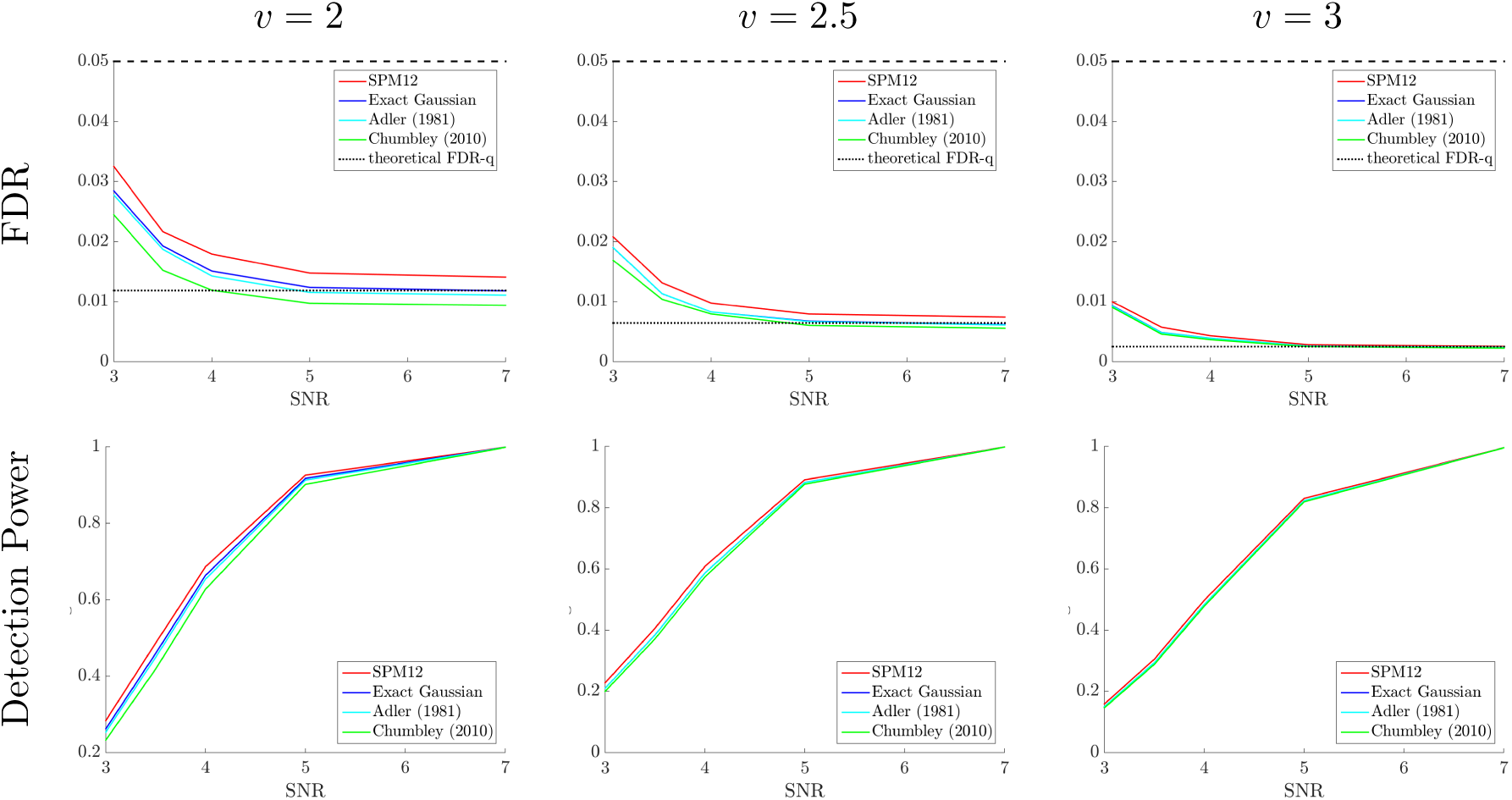
Realized FDR and average detection power (10,000 simulations) as a functionn of pre-threshold. Simulated fields are isotropic Gaussian (Gaussian kernel with bandwidth *ς* = 7).

The third set of simulations considers *t*-fields. The only difference with the Gaussian scenario above is that, for each Monte Carlo simulation, we generate *n*+ 1 independent Gaussian fields *Z*_1_(*s*)*, …, Z _n_*(*s*) and *ε*(*s*) as before, but try to detect peaks of the non-central *t-*fields with *N* d.f.

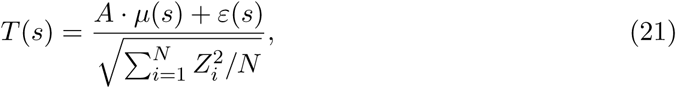

where again *A* is calibrated to achieve the correct SNR for the second highest bump. The construction (21) simulates a situation where the signal is estimated from an average of *N* independent field and detected using a *t* statistic.

Figure 12 compares the realized FDR and detection power, as a function of the number of d.f. of the *t*-field, for various methods. As the number of d.f. increases, the FDR tends to its nominal limit for Gaussian fields. In all cases, using the exact height distribution with no pre-threshold keeps the FDR below the nominal level and results in the highest detection power. This is despite the height distribution being misspecified for *t*-fields.

**Figure 12.**
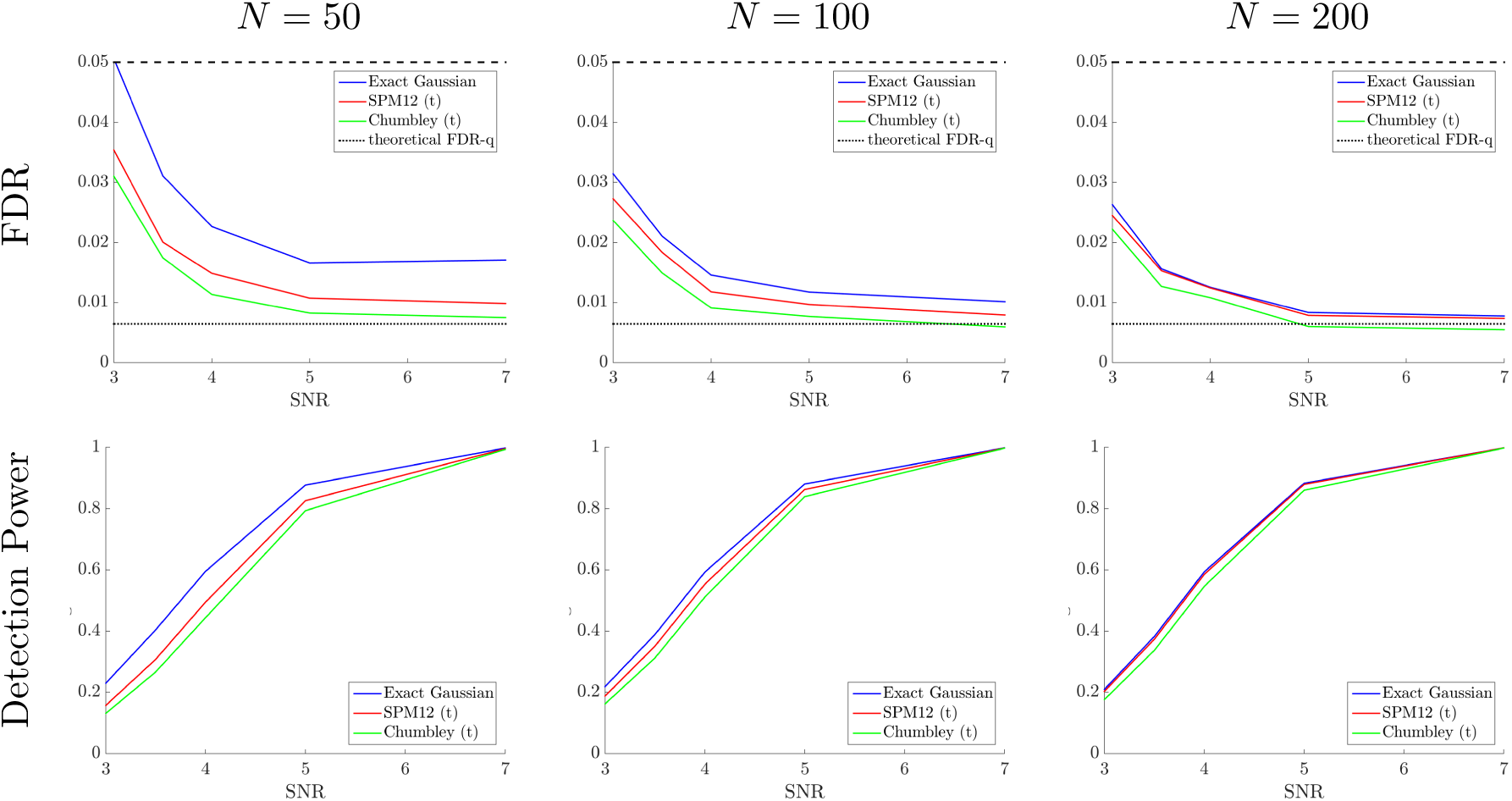
Realized FDR and average detection power (10,000 simulations). Simulated fields are isotropic *t* with *N* d.f., generated from isotropic Gaussian fields (Gaussian kernel with bandwidth *ς* = 7). The pre-threshold is fixed at *v* = 2.5.

## 4 Application to fMRI activation

### 4.1 Voxelwise linear regression

In a single-subject analysis, let

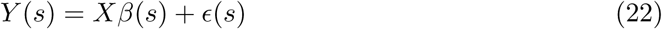

denote the general lineal model (GLM) of the fMRI signal *Y*(*s*) at spatial location *s*, where *X* is the design matrix encoding the task stimulus, drift and other covariates, and *ε(s*) is a noise vector whose entries are assumed to be i.i.d. with zero mean and variance *σ*^2^(*s*). For a fixed vector *c*, the least-squares estimate of the contrast *η*(*s*) = *c*^T^*β*(*s*) is

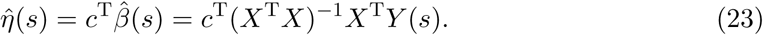

Substituting the model equation (22) and rewriting

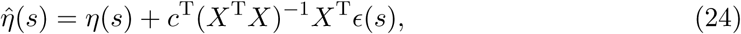

we see that this expression has the same form as the signal+noise model (17), where *η*(*s*) plays the role of the signal, the noise is the field 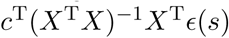, and the observation is the estimated field 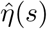. Thus, we may use the methods described above to detect peaks in the parameter surface *η*(*s*). To test the null hypothesis *H*_0_: *η*(*s*) = 0 at each location *s*, the test statistic is the Wald statistic

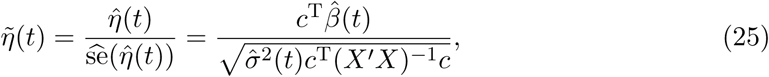

where 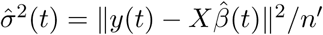 and *n^′^* is the number of degrees of freedom.

Following the prescribed recipe for peak detection, a test statistic map is first produced and the local maxima of this map are found. Each peak is then assigned a p-value and the BH procedure is applied to the list of p-values.

### 4.2 Data example

As a real data example, we use the fMRI data set from Moran et al. (2012), obtained from the public repository OpenfMRI (openfmri.org). Technical details on the full experiment and imaging protocol may be found in the original publication. Among the various experiments conducted in that study, we here focus on the “false-belief task”, whose goal is to find brain regions that are active when processing other people’s false beliefs about reality, in comparison with similar purely physical false realities. Briefly, subjects read short stories corresponding to either a person’s false belief about reality or false realities not involving people. The effect sought after is the contrast between the neural activity in those two states.

As a single-subject analysis, we show here results for subject #49, the same subject analyzed in Cheng and Schwartzman (2017). After pre-procesing, including motion correction and removal of missing data, the scan consisted of 179 time points with image size 71 × 72 × 36 voxels, each voxel measuring 3 mm in all three directions. The data was convolved with a 3D isotropic Gaussian kernel with bandwidth *ς* = 1.6 voxels, corresponding to a FWHM of 3.736 voxels or 11.3 mm. For computational purposes, the kernel was truncated at 2.5 standard deviations from the mode, yielding a kernel support of 8 × 8 × 8 voxels. After convolution, only the valid portion of the image was retained, i.e. those voxels whose values were computed from neighborhood voxels strictly contained in the original image, yielding a valid image of size 64 × 65 × 29 voxels. A linear model was then fitted against the stimulus design matrix and a *t*-statistic map of the form (25) was computed using the appropriate contrast corresponding to the task of interest mentioned above.

Figure 13 shows the thresholded test statistic map using the exact peak height distribution after a *t*-to-Gaussian quantile transformation of the form (10). The significance threshold here is 3.00, equal to the height of the smallest significant peak after BH. Assuming isotropy, the estimate of *κ* for these data is 0.607, yielding the map on the left and containing 31 significant peaks. While the residual fields in these data give the impression of spatial isotropy, justifying the estimation of *κ*, the right panel shows the result when *κ* is fixed and equal to 1, corresponding to a Gaussian autocorrelation function and a noise field that is not necessarily isotropic or stationary. Not surprisingly, forgoing the isotropy assumption yields a more conservative result, with significance threshold 3.75 and 14 significant peaks.

**Figure 13.**
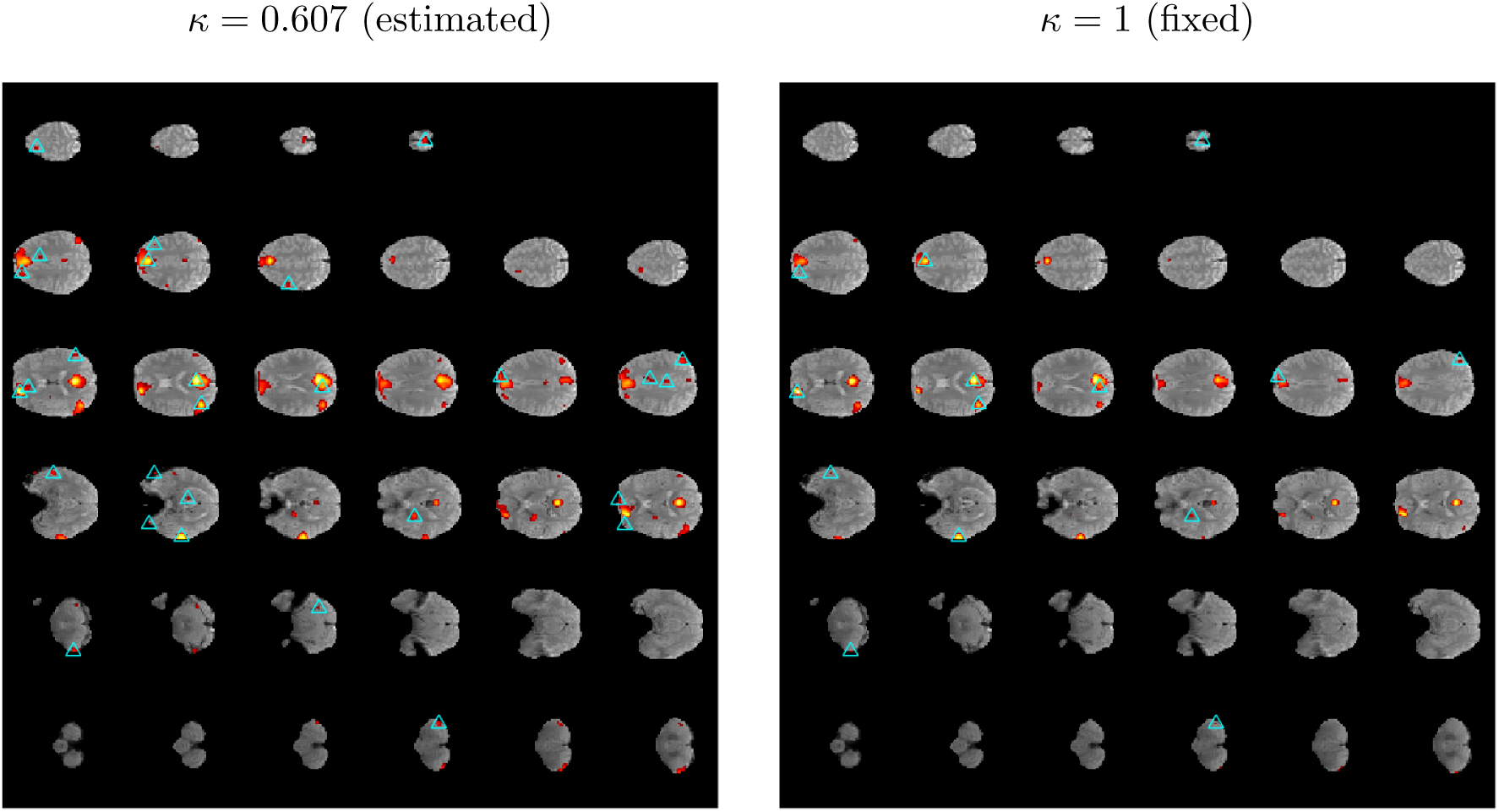
Analysis of the Moran et al. (2012) data: thresholded maps at FDR level 0.05 using the exact height distribution of local maxima, after *t*-to-Gaussian quantile transformation, for isotropic noise with estimated *κ* = 0.607 (left panel) and fixed *κ* = 1 (right panel). In each panel, montage shows the brain volume as transverse slices from the bottom of the brain (bottom left) to the top of the brain (top right). Significant local maxima are marked by cyan triangles. Colored regions indicate the smoothed Wald statistic field above the height of the smallest significant local maximum. Results are superimposed on an anatomical brain image (gray) for reference.

To compare with other methods, Figure 14 shows the activation maps obtained using the overshoot distribution with pre-threshold fixed at *v* = 2.5. The Chumbley and SPM12 p-values were calculated most favorably using their *t* versions with no quantile transformation. In this analysis, the Adler, Chumbley and SPM12 approximations yielded the same activation maps, shown in the right panel as a single image.

**Figure 14.**
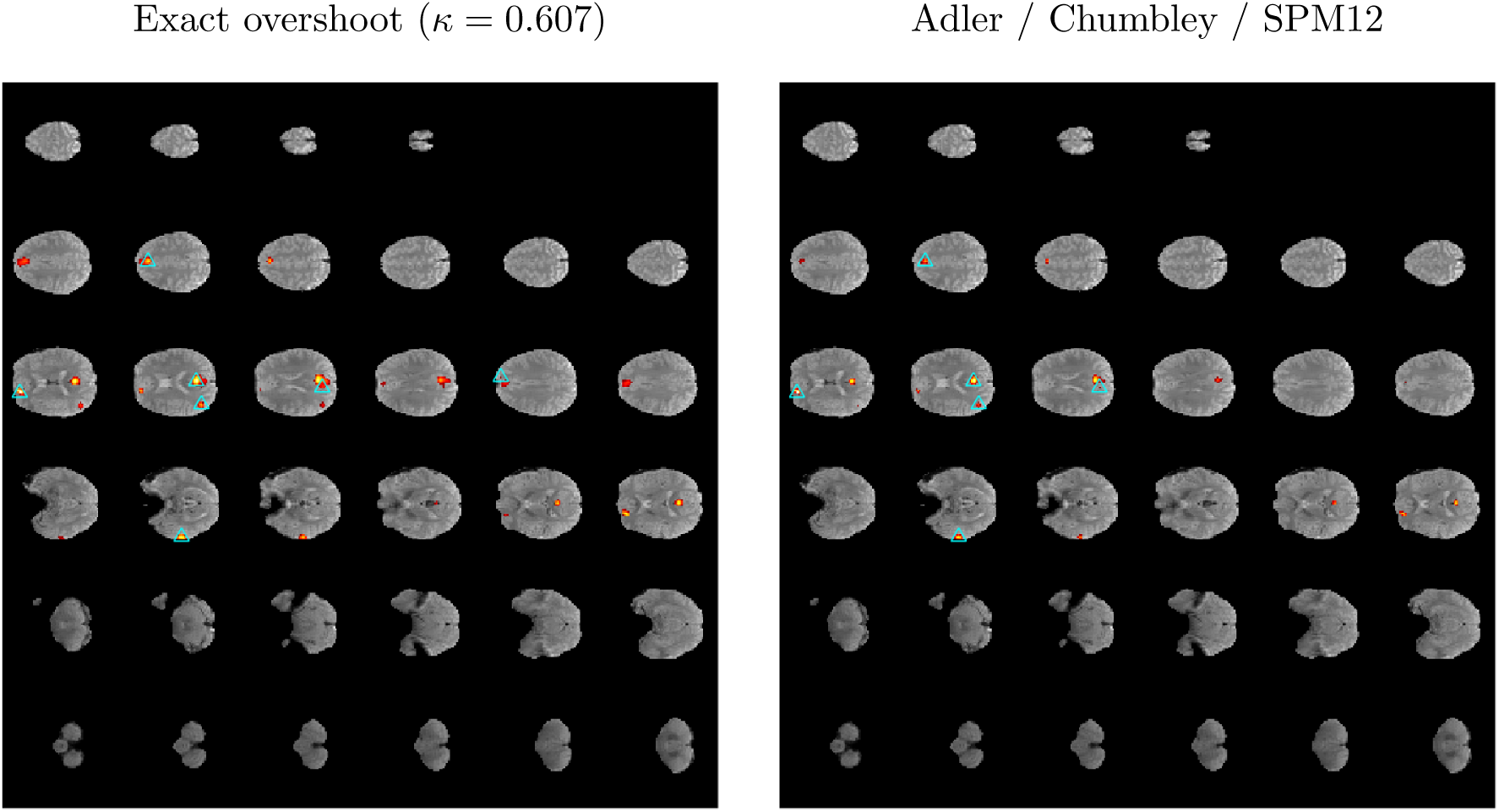
Analysis of the Moran et al. (2012) data: thresholded maps at FDR level 0.05 using the exact overshoot distribution of local maxima, after *t*-to-Gaussian quantile transformation, for isotropic noise with estimated *κ* = 0.607 (left panel) and the approximate overshoot distribution for *t*-fields with no quantile transformation (right panel). The screening pre=threshold is *v* = 2.5. In this particular instance, all three approximate methods yield the same significance threshold and hence the same activation map (right panel). Montages follow the same format as in Figure 13.

Tables 2 and 3 show the significance thresholds and number of significant peaks obtained for various screening pre-thresholds. For every value of the pre-threshold, the exact overshoot distribution with estimated *κ* yields the most significant peaks, followed by fixed *κ* = 1, Adler, Chumbley and finally SPM12. Notice too that applying a pre-threshold and using the overshoot distribution results in a loss of many significant peaks with respect to the exact height distribution with no pre-threshold.

**Table 2.** Analysis of the Moran et al. (2012) data: height significance thresholds for five different methods.

| Distribution | Height | Overshoot |  |  |
| --- | --- | --- | --- | --- |
| | | $v = 2$ | $v = 2.5$ | $v = 3$ |
| Exact, $\kappa = 0.607$ (estimated) | 3.00 | 3.98 | 4.57 | 4.13 |
| Exact, $\kappa = 1$ (fixed) | 3.75 | 4.57 | 4.57 | 4.57 |
| Adler (1981) | - | 4.13 | 4.57 | 4.04 |
| Chumbley et al. (2010) | - | 4.57 | 4.57 | 4.57 |
| SPM12 | - | 4.57 | 4.57 | 4.13 |

**Table 3.** Analysis of the Moran et al. (2012) data: Number of significant peaks for five different methods.

| Distribution | Height | Overshoot |  |  |
| --- | --- | --- | --- | --- |
| | | $v = 2$ | $v = 2.5$ | $v = 3$ |
| Exact, $\kappa = 0.607$ (estimated) | 31 | 9 | 6 | 7 |
| Exact, $\kappa = 1$ (fixed) | 14 | 6 | 6 | 6 |
| Adler (1981) | - | 7 | 6 | 8 |
| Chumbley et al. (2010) | - | 6 | 6 | 6 |
| SPM12 | - | 6 | 6 | 7 |

## 5 Discussion

### 5.1 Peak height vs. overshoot

The main reason for working with the overshoot distribution rather than the peak height distribution is historical. Expressions for the height distribution of isotropic fields were not available in the past. In this paper we have provided an exact formula for the height distribution of peaks, not requiring a pre-threshold. The peak height distribution has been proven theoretically to be correct for isotropic fields and was shown here by simulations to also perform well when the field is anisotropic or slightly non-stationary. The exact height distribution for Gaussian fields is also a good approximation for *t*-fields with high number of d.f., if the *t*-field is transformed to Gaussian by a marginal quantile transformation.

The only practical disadvantage of the exact height distribution is the need to estimate the *κ* parameter. We have seen in the simulations and data analysis above that this is not a problem if one may take advantage of the isotropy of the field to obtain a global estimate. One may also take a conservative value *κ* = 1.

When comparing the exact peak height distribution to the exact overshoot distribution, we have seen that the former results in a higher detection power. The reason is that screening unnecessarily removes potential true peaks to be found. However, the overshoot distribution may still be useful as a conservative approach if the stationarity of the noise field is in doubt. Cheng and Schwartzman (2017) showed that Adler’s approximation is asymptotically valid for high screening pre-thresholds in the case of nonstationary fields.

From the simulation and data analysis results above, the Chumbley and SPM12 approximations appear to be less accurate for Gaussian fields and lead to less powerful inference compared to Adler’s approximation of the exact overshoot distribution. In the case of *t*-fields, the Chumbley approximation is most accurate. In all cases, the SPM12 approximation is the least accurate and least powerful of the methods considered here.

In addition to these results, the approximation currently implemented in SPM12 has the disadvantage that it depends on estimation of the LKCs or resel counts, introducing additional variance and computation time. It should be noted that the marginal p-values reported in the graphical user interface of SMP12 are not peak p-values as required by definition (2) but marginal p-values according to (1). As discussed above, these marginal p-values are misleading low and irrelevant to peak inference.

### 5.2 FDR interpretation

In this paper, we have provided a signal+noise model composed of true unimodal peaks with finite support to be detected against a noisy background. We have argued that peak detection as prescribed by Chumbley et al. (2010) cannot be interpreted as claimed there. For peak FDR to be meaningful, it must be interpreted in the context of the model provided here instead.

Despite this important claim, we emphasize that the purpose of the above argument is not to invalidate the scientific premise that there may be signal everywhere in the brain, and surely not the neurological arguments supporting it. Our goal is only to provide a framework where statistical inference for peaks may be taken to be valid and properly interpreted.

## A Appendix

### A.1 Numerical calculation of the parameter *κ* for a generic kernel

This section describes how to numerically calculate the parameter *κ* in the case of white noise convolved with an isotropic kernel *w*(*s*).

As stated in definition (6), calculation of the parameter *κ* requires the derivatives of the function *ρ*(*r*), which is an expression of the acf *R*(*s*) as a function of 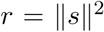. For a field generated as white noise convolved with an isotropic kernel *w*(*s*), the acf *R*(*s*) is the convolution of the kernel with itself. However, computing this convolution analytically and then writing the result as a function of 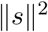 may be difficult. E.g., in the case of the quartic kernel, the convolution is an 8th degree polynomial.

Instead, the following method may be used to calculate the parameter *κ* numerically. First, compute the 3D convolution numerically over a high-resolution grid and extract the values from the center radially in any direction, say, the *x* axis. Matching this vector to the vector *r* = *x*^2^ gives a discrete version of the function *ρ*(*r*). The derivatives *ρ ^′^* and *ρ ^x2033^* are then estimated numerically by differentiation with respect to the vector *r*.

Applying this method to the quartic kernel (7) with *a* = 40 yields the value *κ ≈* 0.92. Recall that *κ* does not depend on scaling (Section 2.1). Choosing a high value of *a* here helps increase smoothness and thus accuracy in the numerical calculations.

## Acknowledgment

This work was partially supported by NIH grant R01 CA157528.

## Footnotes

1 To be precise, 3 times differentiable is not necessary but is sufficient, while 2-times differentiable is necessary but not sufficient. We settle on 3-times differentiable for simplicity.

## References

Adler, R. J., 1981. The geometry of random fields. Vol. 62. SIAM.

Adler, R. J., Taylor, J. E., Worsley, K. J., 2010. Applications of random fields and geometry: Foundations and case studies. URL http://webee.technion.ac.il/people/adler/publications.html

Candes, E., Tao, T., 2007. The Dantzig sselector: statistical estimation when *p* is much larger than *n*. The Annals of Statistics 35 (6), 2313–2351.

Cheng, D., Schwartzman, A., 2015a. Distribution of the height of local maxima of Gaussian random fields. Extremes 18 (2), 213–240.

Cheng, D., Schwartzman, A., 2015b. On the explicit height distribution and expected number of local maxima of isotropic Gaussian random fields. arXiv preprint arXiv:1503.01328.

Cheng, D., Schwartzman, A., 2017. Multiple testing of local maxima for detection of peaks in random fields. Annals of Statistics 45 (2), 529–556.

Cheng, D., Schwartzman, A., 2018. Expected number and height distribution of critical points of smooth isotropic Gaussian random fields. Bernoulli 24 (4B), 3422–3446.

Chumbley, J. R., Friston, K. J., 2009. False discovery rate revisited: FDR and topological inference using Gaussian random fields. Neuroimage 44 (1), 62–70.

Chumbley, J. R., Worsley, K., Flandin, G., Friston, K. J., 2010. Topological FDR for neuroimaging. Neuroimage 49, 3057–3064.

Durnez, J., Degryse, J., Moerkerke, B., Seurinck, R., Sochat, V., Poldrack, R., Nichols, T., 2016. Power and sample size calculations for fMRI studies based on the prevalence of active peaks. bioRxiv, 049429.

Durnez, J., Moerkerke, B., Nichols, T. E., 2014. Post-hoc power estimation for topological inference in fMRI. Neuroimage 49, 45–64.

Efron, B., 2004. Large-scale simultaneous hypothesis testing: The choice of a null hypothesis. J Amer Statist Assoc 99 (465), 96–104.

Eklund, A., Nichols, T. E., Knutsson, H., 2016. Cluster failure: Why fMRI inferences for spatial extent have inflated false-positive rates. Proceedings of the National Academy of Sciences 113 (28), 7900–7905.

Moran, J. M., Jolly, E., Mitchell, J. P., 2012. Social-cognitive deficits in normal aging. J Neurosci 3 (16), 5553–5561.

Pacifico, M. P., Genovese, C., Verdinelli, I., Wasserman, L., 2007. Scan clustering: A false discovery approach. J Multivar Anal 98 (7), 1441–1469.

Poline, J.-B., Worsley, K. J., Evans, A. C., Friston, K. J., 1997. Combining spatial extent and peak intensity to test for activations in functional imaging. Neuroimage 5, 83–96.

Schwartzman, A., Dougherty, R. F., Lee, J., Ghahremani, D., Taylor, J. E., 2009. Empirical null and false discovery rate analysis in neuroimaging. Neuroimage 44 (1), 71–82.

Schwartzman, A., Gavrilov, Y., Adler, R. J., 2011. Multiple testing of local maxima for detection of peaks in 1d. Annals of Statistics 39 (6), 3290–3319.

Worsley, K. J., Marrett, S., Neelin, P., Evans, A. C., 1996. Searching scale space for activation in PET images. Human Brain Mapping 4, 74–90.

Worsley, K. J., Taylor, J. E., Tomaiuolo, F., Lerch, J., 2004. Unified univariate and multivariate random field theory. Neuroimage 23, S189–195.

Zhang, C.-H., 2010. Nearly unbiased variable selection under minimax concave penalty. The Annals of Statistics 38 (2), 894–942.

Zhang, H., Nichols, T. E., Johnson, T. D., 2009. Cluster mass inference via random field theory. Neuroimage 44, 51–61.

